# Identification of the rice *Rc* gene as a main regulator of seed survival under dry storage conditions

**DOI:** 10.1101/2022.12.08.519629

**Authors:** C.T. Manjunath Prasad, Jan Kodde, Gerco C. Angenent, Fiona R. Hay, Kenneth L. McNally, Steven P.C. Groot

**Author notes:** **Author for Correspondence:** Steven P.C. Groot, **email:**.

## Abstract

Seed deterioration during storage results in poor germination, reduced seed vigor, and non-uniform seedling emergence. The rate of aging depends on storage conditions (RH, temperature, and oxygen) and genetic factors. This study aims to identify these genetic factors determining the longevity of rice seeds stored under experimental aging conditions mimicking long-term dry storage. Genetic variation for tolerance to aging was studied in 300 *Indica* rice accessions and storing dry seeds under elevated partial pressure of oxygen (EPPO) condition, using a genome-wide association study. The association analysis yielded eleven unique regions across the genome for all measured germination parameters after aging. These genomic regions differed from regions previously identified in rice under humid experimental aging conditions. The significant single nucleotide polymorphism in the most prominent region was located within the *Rc* gene, encoding a bHLH transcription factor. Storage experiments using isogenic rice lines (*SD7-1D* (*Rc*) and *SD7-1d* (*rc*)) with the same allelic variation confirmed the functional role of the *Rc* gene, conferring a stronger tolerance to dry EPPO aging. A functional *Rc* gene results in the accumulation of pro-anthocyanidins in the pericarp of rice seeds, an important sub-class of flavonoids having strong antioxidant activity, which may explain why genotypes with an allelic variation for this gene show variation in seed tolerance to dry EPPO aging.

## INTRODUCTION

The use of good-quality seeds is the foundation of global food security. Seed longevity, a vital component of seed quality, is of paramount importance for the seed industry, farmers, genebanks, and restoration of terrestrial ecosystems. Seed longevity (or storability) is defined as the capacity of the seeds to germinate after storage (Barton, 1961; Justice and Bass, 1978). During storage, deterioration progresses with time. Seed survival after the storage is a result of a complex interplay between initial seed quality, storage conditions (relative humidity (RH) or seed moisture content, temperature, and oxygen), and genetic makeup (McDonald, 1999). A major cause of seed deterioration is free-radical (reactive oxygen species, ROS) mediated damage to macro-molecules and bio-membranes (Bailly, 2004; Halliwell and Gutteridge, 2015; Fleming et al., 2018; Waterworth et al., 2019). To resist damage, seeds accumulate protective substances such as LEA proteins, enzymes, non-reducing sugars, and antioxidants during their development (Bentsink et al., 2000; Sattler et al., 2004; Kalemba and Pukacka, 2007; de Souza Vidigal et al., 2016; Leprince et al., 2016; Petla et al., 2016; Lee et al., 2017). The protective mechanisms that operate depend on the moisture status of the seed; for example, scavenging of ROS in the dry seed is largely due to antioxidant molecules such as tocopherols and glutathione, while enzymatic antioxidants such as catalases become active upon moistening of the seeds (Gerna et al., 2022).

Storing seeds under proper conditions is essential for preserving high seed quality, vigor, and viability (Priestley, 1986; Bewley and Black, 1994; Hong et al., 1996). Commercial seed lots of many vegetables (low-volume and high-value seeds) are often stored under controlled environments (30% RH and 15 or 20°C) to maintain high seed quality. Unlike vegetable seeds, many cereal crop seeds (high-volume and low-value seeds), including rice seeds, are often dried to 12% moisture content followed by storage in warehouses under uncontrolled conditions. Further, packing seeds in moisture-permeable bags allows moisture exchange. Consequently, uncontrolled conditions under humid tropical climates result in a rapid loss in seed quality in most rice seed storage facilities and hence germination may drop below 70% within six months. The seeds that survive will germinate with low vigor and give rise to relatively weaker seedlings with slower root growth and poor seedling establishment than those obtained from fresh (non-stored) seeds.

Rice (*Oryza sativa* L.) is the second most important cereal crop, and the main staple food for more than half of the world’s population (Chauhan et al., 2017). Sustainable rice production is challenged by the need for dry cultivation and mechanical transplanting or direct seeding to reduce water use and cope with labor shortages (Mahender et al., 2015). The use of high-quality, better germinating seeds will be an essential part of the answer. The area for hybrid rice is expanding, and, since hybrid seeds are more expensive to produce, efforts to prolong their shelf-life are needed. Considering the high seed volume and costs involved, rice seeds cannot be stored in the tropics under cooled conditions like vegetable seeds. Thus, dry storage could be an option for commercial rice seed storage. Further, breeding for cultivars with higher seed longevity has the potential to maintain seed quality longer during storage.

Understanding seed longevity traits and their underlying genetic determinants remains a scientific challenge (Righetti et al., 2015; Sano et al., 2016; Liu et al., 2022). Genetic analyses for seed longevity in rice have identified several QTLs in mapping populations derived from crosses between Japonica and Indica variety groups (Miura et al., 2002; Sasaki et al., 2005; Zeng et al., 2006; Xue et al., 2008; Jiang et al., 2011; Hang et al., 2015; Lin et al., 2015; Dong et al., 2017; Yuan et al., 2019). These QTL studies revealed that alleles from Indica varieties promoted seed longevity in each population. A major QTL on chromosome 9 is considered the most reliable and stable QTL related to seed longevity (Shigemune et al., 2008; Li et al., 2012). Fine mapping of QTL *qLG-9* using advanced backcross progeny identified a potential candidate gene (*TPP7* encoding a trehalose-6-phosphate phosphatase) for seed longevity (Sasaki et al., 2015). Since longevity experiments under natural dry conditions take a long time, longevity parameters are often derived from experimental aging tests like controlled deterioration (CD; (Powell, 1995)) or Artificial Aging (AA; TeKrony (1993)), where seeds are placed under relatively high humidity and temperature conditions to speed up deterioration (Hay et al., 2019). However, the results of humid aging tests often show a poor correlation with long-term seed storage under dry conditions (Schwember and Bradford, 2010; Agacka-Mołdoch et al., 2016), including rice seeds (Hay et al., 2019). One apparent reason is that in moist seeds, enzymes such as catalases can be active, which is not the case at low water activity levels (Barbosa-Cánovas et al., 2020).

Seed aging under dry ambient conditions can be accelerated by storing at higher oxygen concentrations (Roberts, 1961; Schwember and Bradford, 2011; Groot et al., 2015; Gerna et al., 2022) and elevated air pressures (Groot et al., 2012; Nagel et al., 2016; Hourston et al., 2020; Renard et al., 2020). Oxygen and ROS are detrimental to seeds during storage, e.g., by inducing damage to genetic material (Moutschen-Dahmen et al., 1959; Ohlrogge and Kernan, 1982), reducing tocopherol levels and impairing mitochondrial functioning (Groot et al., 2012). Genetic studies with barley and *Arabidopsis* have identified several QTLs for longevity under dry experimental aging conditions using elevated partial pressure of oxygen (EPPO), which differ at least partly from those identified under more moist storage conditions (Nagel et al., 2016; Buijs et al., 2020; Renard et al., 2020). At the genetic level, EPPO storage also mimics dry after-ripening to release dormancy in *Arabidopsis* seeds, which identified *DOG* loci also previously described for dormancy release by after-ripening during long-term laboratory bench storage (Buijs et al., 2018). Recently, a genome-wide association study (GWAS) using a diverse Indica rice panel has identified eight major genomic regions for seed longevity parameters measured from seeds stored relatively dry (60% RH) at 45°C (Lee et al., 2019). In that study, the candidate genes responsible for increased longevity are involved in mechanisms related to DNA repair, transcription regulation, ROS scavenging, and embryo/root development.

Here we report on a GWAS in rice using diverse *Indica* rice accessions aiming to study the genetic variation in seed survival under dry (50% RH) EPPO experimental aging conditions compared with ambient control conditions. Our experiments aimed to: (1) assess natural genetic variation for seed germination and associated parameters following dry-EPPO storage, (2) identify potential candidate genes associated with seed longevity through GWAS, and (3) validate the functional role of a significant candidate gene conferring tolerance to dry-EPPO storage conditions.

## RESULTS

### Optimum dry aging duration for phenotyping the GWAS population

A pre-test with seed samples from 20 accessions showed an apparent decline in average seed germination after the EPPO treatments, whereas there was a small but significant decline as a result of the EPPN (pressure control) treatments and no significant decline in the ambient control during storage up to 56 days (Figure 1). The limited number of samples used in this pre-test does not allow for verification of whether the observed germination response is only a genotype effect or also because of seed production conditions. However, for simplicity, we consider here the germination response as an accession effect. The descriptive statistics and ANOVA results, showing variation in accessions (A), storage durations (S), and their interactions (AxS) under different aging treatments (T), are detailed in Supplemental Table 2. Among all three aging treatments, a significant difference in total seed germination was observed among seed samples from different accessions (Supplemental Table 2). Accession samples stored for 21 days in the EPPO aging treatment recorded average total seed germination of 47% with a range between 9 and 91%. After 42 days EPPO storage, the average total seed germination was below 10%, and at 56 days, seeds of all accessions showed a complete loss in viability. Under this treatment, variation in total seed germination differed significantly between accession samples (A; *p*<0.001), storage duration (S; *p*<0.001), and their interaction (AxS; *p*<0.001). Based on the observed variation, 21 days of storage was chosen to screen the GWAS population for different seed germination parameters.

**Figure 1.**
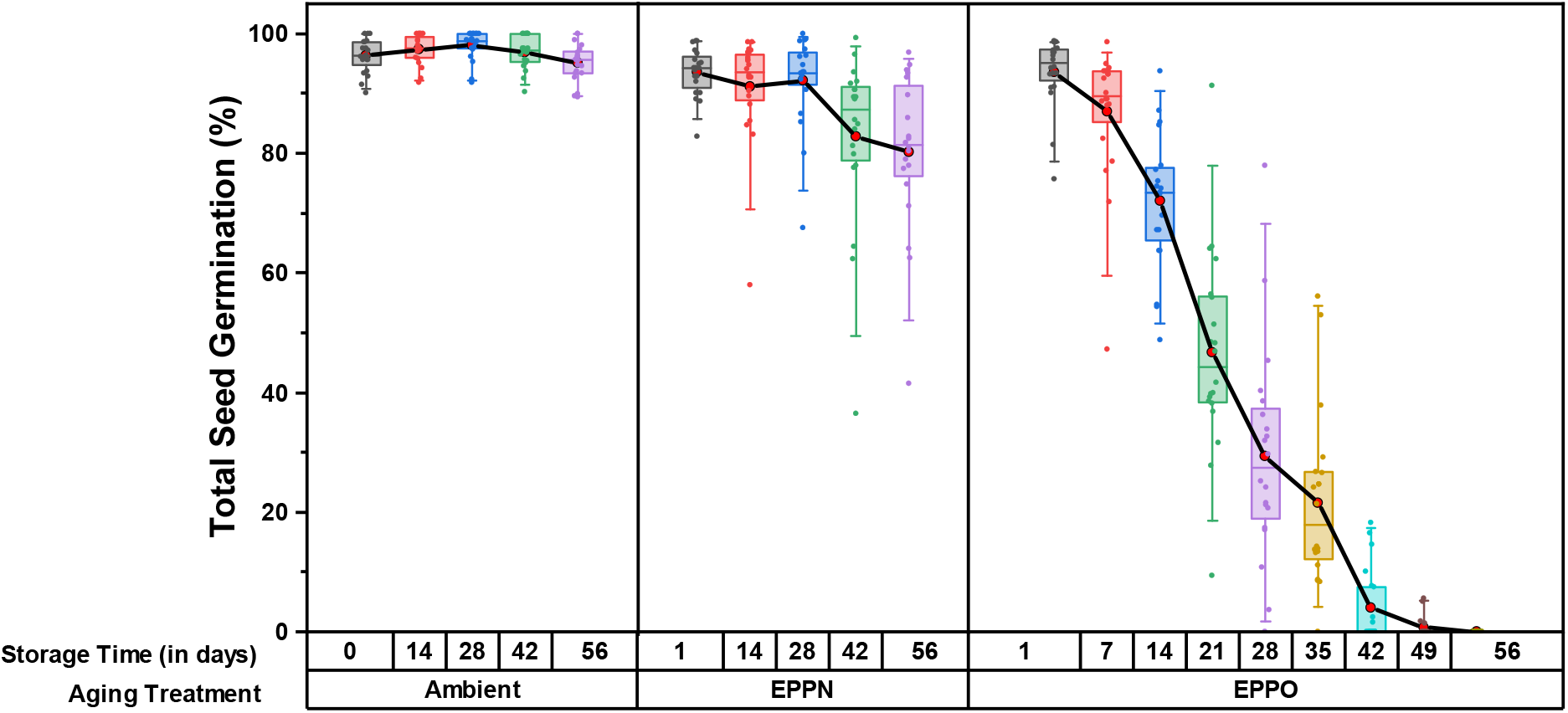
Phenotypic variation for total seed germination (%) among 20 rice accessions under different aging treatments used in the pre-test. Box plot showing distribution for average total seed germination over different storage time. Aging treatments were performed on seeds at 50% RH and 35°C (n=2 × 40-45 seeds).

### The GWAS panel shows large variation in germination parameters in response to dry aging treatments

In the main test, 21 days storage of seed samples from the 300 accessions under EPPO aging treatment showed considerable variation for all the germination parameters measured (Figure 2 & Supplemental Table 3), with significance for accessions (A; *p*<0.001), aging treatments (T; *p*<0.001) and their interactions (AxT; *p*<0.001) (Supplemental Table 3 & 4). Total seed germination ranged from 5 to 97% after 21 days EPPO aging treatment, from 43 to 100% after 21 days EPPN pressure control, and from 84 to 100% for ambient control (0 and 21 days) treatments (Figure 2A & Supplemental Figure 3). The average total seed germination was 50% after 21 days EPPO aging treatment, which was significantly lower than the EPPN pressure control (89%) and ambient controls (97%) (Figure 2A). The slower and less total germination resulted in a lower average value for area under the germination curve (AUC(250)) after EPPO aging was 52 compared with 161 for the 21 days EPPN pressure control treatment, and 191 for the ambient 0 and 21 days control treatments (Supplemental Figure 4A). The average number of normal seedlings was 23% after EPPO 21 DOS aging treatment (Supplemental Figure 4B & Supplemental Figure 5). The ΔEPPO trait values, corrected for the pressure control (EPPN), also showed large variation for all the germination parameters and were significantly different from the 21 days EPPO and control aging treatments. After this correction (ΔEPPO), average total seed germination, area under the germination curve (AUC(250)), and total normal seedlings were respectively 58%, 81.3 h, and 32% after 21 DOS, which were significantly lower than those for the 0 and 21 days ambient pressure control treatments (Supplemental Table 3). The Shapiro-Wilk normality test showed an improved normality of transformed trait values for GMAX when compared to the original trait values (Supplemental Figure 6). The observed large variation for GMAX under each aging treatment was mainly attributed to the accession effect, with only a minimal effect of different production years (2009-2016) (Supplemental Figure 7).

**Figure 2.**
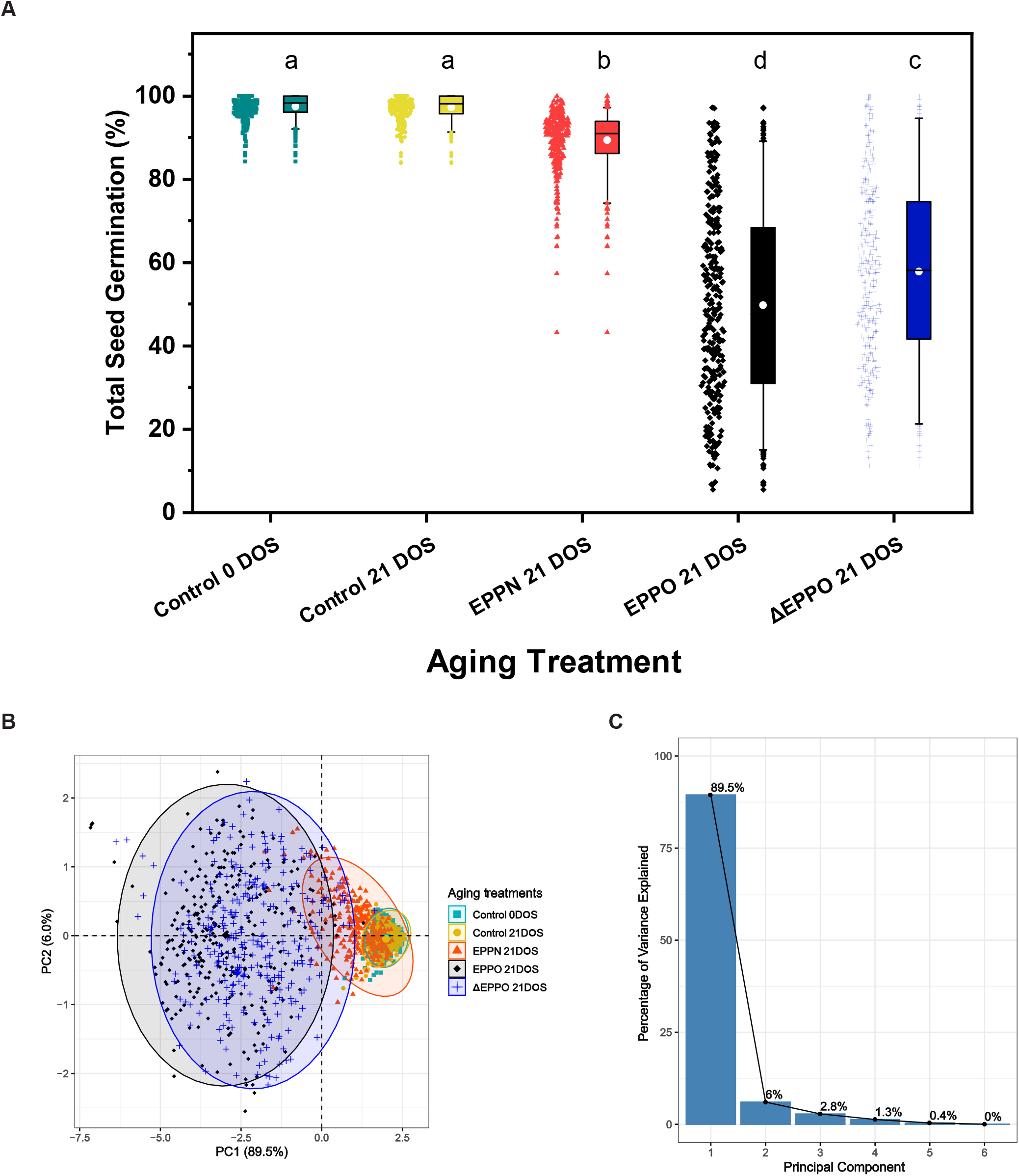
Phenotypic variation observed among 300 rice accessions for total seed germination under different aging treatments in the main test. **A**. Box plot showing distribution for total seed germination. Aging treatments includes Control 0 days of storage (DOS) (■), Control 21 DOS (●), EPPN 21 DOS (▲), EPPO 21 DOS (♦) and ΔEPPO 21 DOS (+). Aging treatment was performed on seeds at 50% RH and 35°C for 21days (n= 2×40-45 seeds). **B-C**. Principal component analysis (PCA) of different seed germination parameters. PCA plot for first two PCs under different aging treatments **(B)** and scree plot **(C)** indicating percentage of phenotypic variance explained by each principal component.

The principal component analysis (PCA) revealed variations among the germination parameters under different aging treatments (Figure 2B & Supplemental Table 5). The first two principal components (PC1 and PC2) cumulatively explained 95.5% of total phenotypic variation across aging treatments (Figure 2C). Correlation coefficient analysis showed the existence of a significant positive correlation (r = 0.94) between the total seed germination data recorded in the pre-test and the main test (Supplemental Figure 8). Here, we focus on the genetic association for total seed germination (GMAX).

### Population genetic structure and whole-genome linkage disequilibrium

The population structure or genetic relatedness should be accounted for in GWAS to avoid the identification of spurious associations (Sul et al., 2018). Increasing *K* from 4 to 5 in the ADMIXTURE analysis still decreased the CV error substantially (Supplemental Figure 9A), whereas at *K*>5 there is no substantial decrease in the CV error estimate, suggesting *K*=5 as the most likely number of clusters. The population subdivision at *K*=2 reproduced the first PCA coordinate by separating the Xian/Indica subpopulation 1A (XI-1A) (Wang et al., 2018) from all other groups (Supplemental Figure 9B). The principal component analysis (PCA) based on the DNA sequence data showed a clustering of accessions into five groups (Supplemental Figure 10). The first three principal components (PCs) explain 15.32% of the total genetic variation (Supplemental Figures 10A & 10B). The number of rice accessions grouped into different sub-groups is illustrated in Supplemental Figure 10C. The extent of LD and its decay with genetic distance is critical for determining the number of markers (SNPs) needed to successfully map a QTL related to the phenotype. The LD decay pattern in the total population used in the experiment was estimated based on LD squared correlation coefficient (*r*^*2*^) between pairs of SNPs. The average LD across chromosomes dropped to half of its initial value at ∼180 Kb (Supplemental Figure 10D). Therefore, we choose to consider the 200 Kb distance on either side of the most significant SNP as a single genomic region harboring potential candidate genes.

### GWAS identifies genomic regions associated with total seed germination across aging treatments

Following phenotypic characterization of 300 accessions, genome-wide association analysis was performed with probit-transformed trait values to identify sets of SNPs statistically associated with total seed germination across aging treatments (Supplementary Table 6). In total, 14 significant genomic regions were identified across the aging treatments (Figure 3); the details of each region are presented in Supplemental Table 7. Two regions were identified in relation to the ambient control pre-storage treatment (0 DOS), one to ambient control storage for three weeks (21 DOS), three to pressure control treatment (EPPN), and four each to elevated oxygen pressure treatment (EPPO) and corrected EPPO treatment (ΔEPPO) (Figure 3 & Supplemental Table 7). Using the 200 Kb borders around the most significant SNPs, there were no common regions between controls and EPPO aging treatments, indicating that in our experiment, sensitivity to the storage under ambient or pressure control (EPPN) treatments are genetically independent of storage under an elevated partial pressure of oxygen (EPPO). We observed a common genomic region on chromosome 7 (R7 and R9) between EPPO and ΔEPPO treatments (Figure 3D & 3E). In pressure control treatment (EPPN 21 DOS), the less frequent allele of region R4 had a negative effect, whereas less frequent alleles in the other regions showed a positive effect (Supplemental Table 7). In general, the strongest allelic effects (AE) were observed in the EPPO treatment (Supplemental Table 7). The most significant and prominent genomic region identified in EPPO 21 DOS and ΔEPPO 21 DOS was ‘R7’, with a lowest -log_10_(*p*) value and a strong allele effect of >33% (Supplemental Table 7). Thus, the potential role of one or more candidate genes present in this region are of prime importance in identifying their role in seed longevity under dry storage.

**Figure 3.**
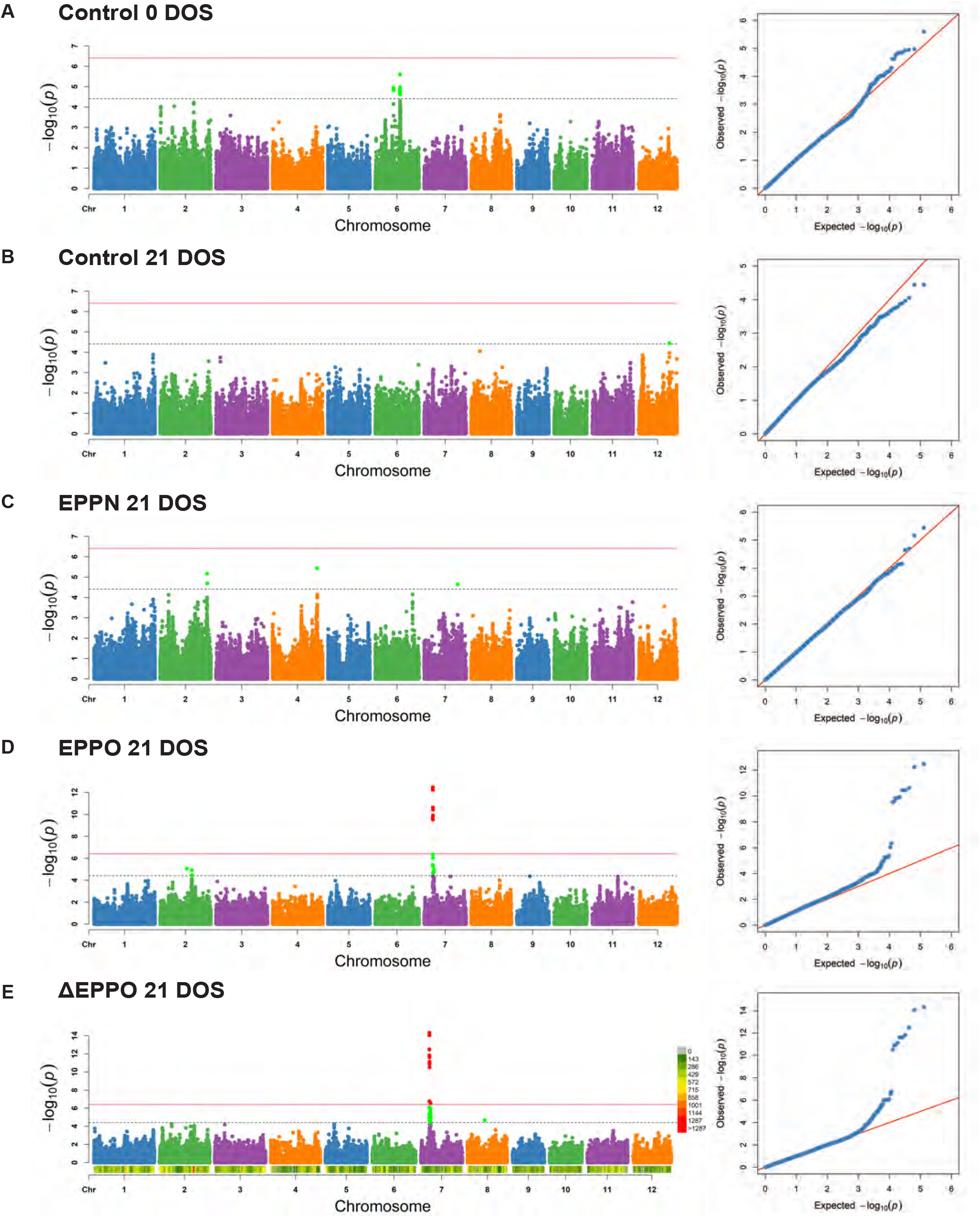
Genome-wide association analysis for total seed germination under different aging treatments. **A-E**. GWAS results for total seed germination for different aging treatments-Control 0 days of storage (DOS) **(A)**, Control 21 DOS **(B)**, EPPN 21 DOS **(C)**, EPPO 21 DOS **(D)** and ΔEPPO 21 DOS **(E)**. GWA analysis was performed using 1M GWAS SNP dataset on probit-transformed values. Manhattan plot to the left indicate SNPs from each chromosome along x-axis and the -log_10_(*p*) values for the association along the y-axis. Significant SNPs are colored red if the -log_10_(*p*) value is greater than the Bonferroni corrected threshold (solid red line) and colored light green if the -log_10_(*p*) value is between -log_10_(*p*) value = 4.41 threshold (dotted black line) and Bonferroni corrected threshold. Quartile-Quartile (QQ) plot to the right indicate the expected versus the observed -log_10_(*p*) values.

### Candidate genes underlying the genomic regions for total seed germination across aging treatments

The LD decay rate of the population used in our study was around 200 Kb (Supplemental Figure 10D). However, the resolution of significant associations at each genomic region varied due to the local LD patterns. For each region, we determined LD blocks harboring an identified significant SNP in a region containing candidate genes. A detailed description of these regions, including gene annotations and functions, are presented in Supplemental Table 8. In most cases, the identified significant SNPs of the genomic regions were either within or close to a gene. Since our experiments mainly aimed to unravel the genetics behind the observed phenotypic variation in sensitivity or tolerance of seeds to aging during dry storage, we focus here on the putative candidate genes in the genomic regions significantly associated with ΔEPPO.

A total of 10 candidate genes from 4 different genomic regions was identified for the trait values recorded under ΔEPPO 21 DOS. The prominent genomic region, ‘R7’, was also identified with original or non-corrected values of EPPO aging treatment (Supplemental Table 7). Candidate genes in region R7 on chromosome 7 explained the variation observed for total seed germination. The most significant SNP at region R7 was positioned within the *OsRc* gene.

For total seed germination, the significant SNP on region R1 in chromosome 6 was close to a gene coding for *Cytochrome P450* (LOC_Os06g30500.1), a protein that exhibits oxidoreductase activity and represents a family of haem-containing enzymes involved in synthesis or degradation of several hormones including GA, auxin, and brassinosteroids. Likewise, two candidate genes in region R2 on chromosome 6, *Transducing Family Protein* (LOC_Os06g22550.1) and *Formyl Transferases* (LOC_Os06g22560.1), are involved in nucleotide binding activity and purine biosynthesis, respectively. Similarly, for genomic regions (R4 to R6) associated with trait values recorded for seeds aged under high-pressure nitrogen (EPPN 21 DOS), seven candidate genes were identified as mainly having a role in the regulation of plant defense and transcription.

Since the major effect genomic region, ‘R7’ (*OsRc*) was common and consistently found in non-corrected and corrected EPPO aging treatment for total seed germination trait values, we further focused on the role of this candidate gene in seed longevity.

### Rc gene in genomic region ‘R7’ has a strong effect on tolerance to oxygen aging

Our search for putative candidate genes in the genomic region ‘R7’ pointed to three candidate genes (Supplemental Table 9, see *R7* and Figure 4A) within the LD block spanning 58 Kb, while the most significant SNP (position=218399762 corresponding to Chr7:6067855) was located within the *Rc* gene (LOC_Os07g11020.1). This gene encodes a basic helix-loop-helix (bHLH) family transcription factor, regulating pro-anthocyanidin (PA) synthesis in the pericarp (Sweeney et al., 2006). Indeed, seeds from accessions with an A allele (n=37) at this SNP had a red-colored pericarp, while accessions with G allele (n=259) showed a light-colored pericarp. Rice accessions with the A allele recorded, on average, a significantly higher germination of 79% compared to accessions with G allele with 54% average germination after 21 days EPPO storage (Figure 4B).

**Figure 4.**
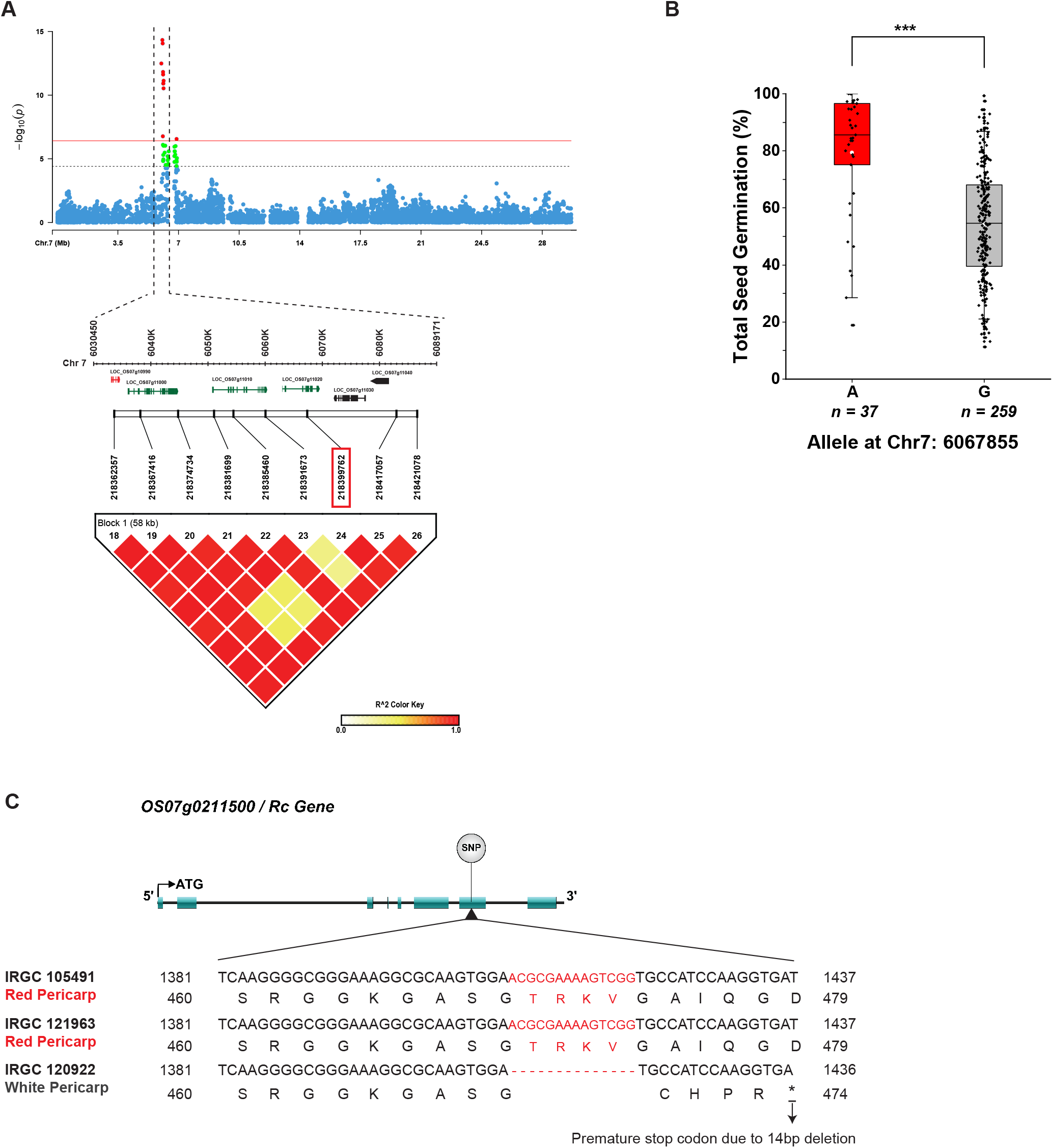
Genomic analysis of the common significant SNP identified for seed germination parameters under ΔEPPO aging treatment. **A**. Manhattan plot showing genome-wide association results for total seed germination along with LD block based on *r*^*2*^ value between SNPs on chromosome-7. The color intensity corresponds to the *r*^*2*^ value according to the legend at the bottom right. The significant SNP (218399762) marked in the red rectangle. **B**. Allelic effect of chromosome-7 locus (LOC_Os07g11020) for percentage total seed germination. Paired t-test shows significant allelic effect difference with reference to major and minor allele. **C**. Multiple sequence alignment showing coding sequence and protein sequence for the gene harboring the most significant SNP specific for EPPO aging treatment identified in the genome-wide association analysis. Sequence comparison is between *Oryza rufipogon* (IRGC 105491) with red pericarp, one rice accession with A allele (IRGC 121963) and one with G allele (IRGC 120922) at the most significant SNP on chromosome-7 identified in this study.

In addition, the full-length coding sequence and protein sequence of the *Rc* gene with an allelic difference were compared along with those of *Oryza rufipogon* (IRGC 105491), showing seeds with a red pericarp color. All accessions with the G allele had a 14bp deletion in the exon-7 of the *Rc* gene, causing a pre-mature stop codon resulting in knocking out its gene function (Figure 4C and Supplemental Figure 10). This functional nucleotide polymorphism explains the change in seed pericarp color between the allele variants.

### Functional Rc gene influences dormancy and longevity phenotypes

To confirm the role of the *Rc* gene in seed longevity, we analyzed seeds from isogenic rice lines *SD7-1D* and *SD7-1d. SD7-1D* was previously identified as an allele influencing rice seed dormancy and is presently known to be identical with the *Rc* gene responsible for a red pericarp (Gu et al., 2011). Here we use *OsRc, Rc* or *rc* when referring to the gene and *SD7-1D* or *SD7-1d* when referring to the isogenic lines having dormancy promoting or decreasing alleles, respectively. The isogenic line *SD7-1D* has a functional *Rc* gene introduced from the weedy red rice line, SS18-2, into the EM93-1 genetic background through single-plant marker-assisted selection and recurrent backcrossing (Gu et al., 2011). The coding sequence of the isogenic lines also differs in the 14 bp deletion in the exon 7 at position 5177-5190, similar to the difference between the A and G alleles in our GWAS population (Supplemental Figure 12), resulting in an early stop codon, which accounts for the lack of a red pericarp color (Figure 5A). When grown for seed production, isogenic lines were similar in plant architecture and other seed morphology traits (Supplemental Figure 13). The mature seeds of isogenic lines clearly showed variation in both dormancy and longevity phenotypes. Under ambient storage conditions (30% RH and 20°C), dormancy release (after-ripening) was slower in seeds with a functional *Rc* allele (red pericarp) when compared to seeds with the *rc* alleles (Figure 5B). Dormancy of fresh seeds of both the isogenic lines was lost in about five weeks under ambient storage conditions (Figure 5B) and within seven days with a dormancy-breaking treatment (Supplemental Figure 14). Interestingly, dormancy was lost more rapidly for seeds from both lines when stored in the EPPO aging treatment compared with ambient storage conditions (Figure 5C). Furthermore, *Rc* seeds from the isogenic line with a red pericarp (*SD7-1D*) showed higher tolerance under EPPO aging treatment compared to *rc* seeds from line *SD7-1d*. Under ambient control and pressure control (EPPN) aging treatments, seeds of both isogenic lines recorded >98% germination at the end of the test (Supplemental Figure 15 and 16). The time to reach 50% germination (t50, in h) after storage in the EPPN pressure control treatment indicates that three days of storage already gives a significant delay in germination for the light-colored pericarp seeds, with an increase in t50 values but with no significant change over storage time. This EPPO effect was less present in the isogenic line with a red pericarp. High total seed germination was retained up to seven weeks of EPPO storage with red pericarp seeds, whereas total seed germination started to decline after three weeks with the light-colored pericarp seeds (Figure 5C). Once viability started to decline, the subsequent decline in total seed germination was faster (the slope of the curve is steep) with *Rc* seeds (red pericarp) when compared with *rc* seeds (light-colored pericarp) that had a more gradual decline over time.

**Figure 5.**
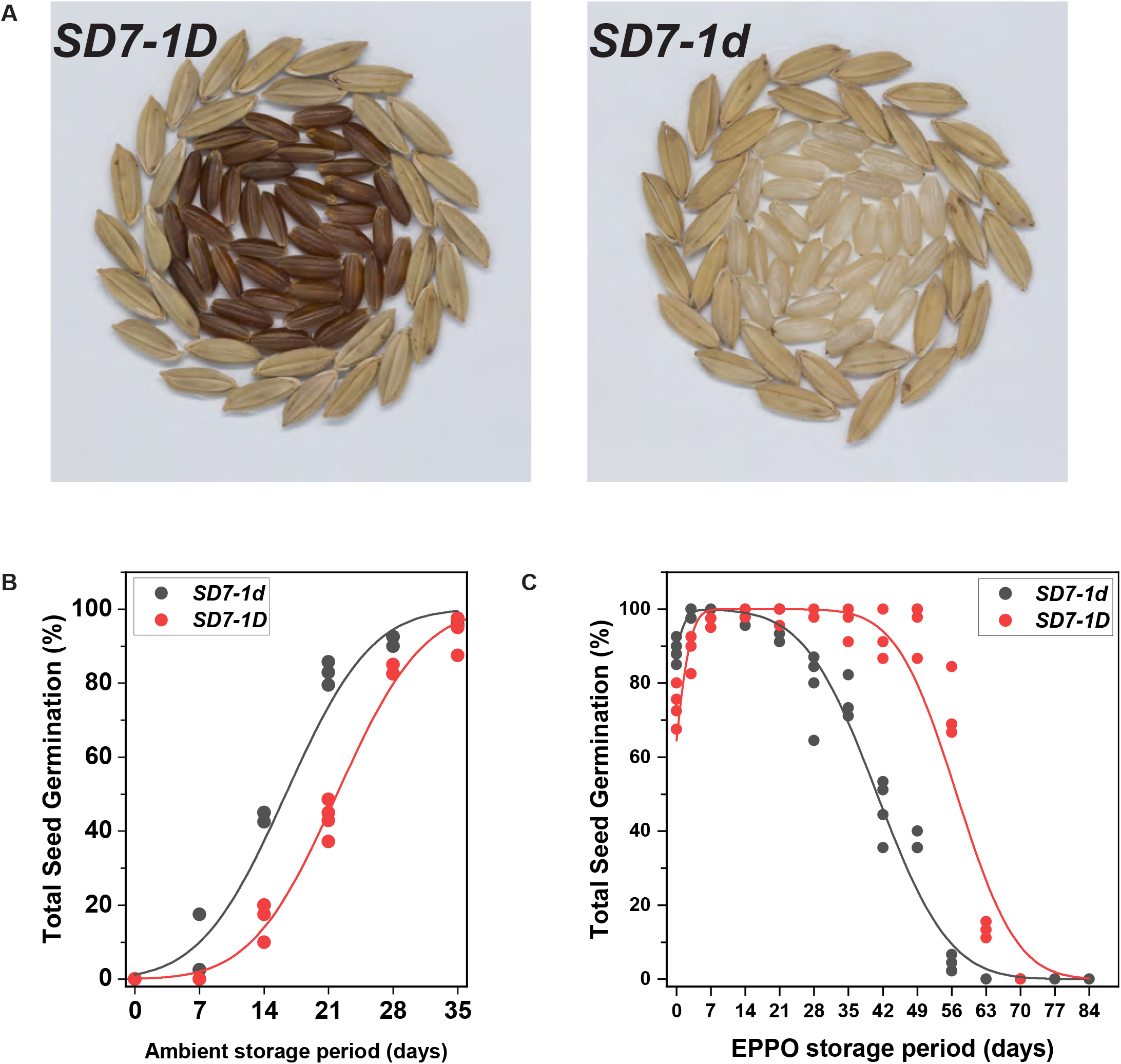
Functional analysis of *Rc* gene (LOC_Os07g11020) identified in GWAS for role in protection of seed during storage under EPPO aging conditions. **A**. Difference for pericarp color of mature seeds between isogenic lines *SD7-1D* with red pericarp and *SD7-1d* with light colored pericarp. Seeds in the outer two rows show seeds with palea and lemma, and the inner rows show dehulled seed or caryopsis. **B**. Comparison of dormancy release (after-ripening) from freshly harvested seeds during ambient storage (30% RH and 20°C) between isogenic lines. Each dot in the plot indicate average values of germination test evaluated on four biological replicates of 45 seeds each at 14 days after imbibition (DAI). **C**. Comparison of germination of isogenic lines under EPPO aging conditions. Freshly harvested seeds equilibrated to 40% seed RH at 20°C for 22days were used. EPPO aging was performed at 50% RH and 35°C for different storage periods. Each dot in the plot indicates average values of germination test evaluated on four biological replicates of 45 seeds each at 14 DAI.

### Analysis of mitochondrial quality

Rice seed samples subjected to different aging treatments were analyzed for ethanol production during the first day after the start of imbibition, a marker for anaerobic respiration related to mitochondrial damage (Supplemental Figure 17). Seeds that had been stored under ambient and pressure control treatments until 21 days produced only a small amount of ethanol. EPPO-aged seeds, in contrast, produced relatively high ethanol levels. In this, seven days EPPO stored seeds with a light-colored pericarp (from line *SD7-1d*) produced more ethanol when compared with the red pericarp seeds (from line *SD7-1D*) (Supplemental Figure 17A). Not much difference was observed between headspace ethanol production from seeds of both isogenic lines stored for 14 days or 21days under EPPO aging treatment (Supplemental Figures 17B and 17C).

## DISCUSSION

Seed lots with higher longevity characteristics will provide better crop establishment after storage. Hence, cultivars with improved longevity have the potential to realize higher yields. Genetic markers conferring improvement in seed longevity would be valuable for breeders, seed companies, and genebank managers. Research on seed longevity is hindered because aging under conventional storage conditions takes too long for selection and due to the lack of convenient aging assays to determine storage potential under dry storage conditions. Deterioration mechanisms involved in moist experimental aging, as in a Controlled Deterioration (CD) test, differ at least partly from those during commercial dry storage (Walters, 1998; McDonald, 1999). As an alternative to moist aging, we used here dry-EPPO aging, where seed deterioration is accelerated using elevated oxygen pressure while the seed moisture levels resemble traditional dry seed storage conditions (60% RH and 28°C) (Prasad C. T. et al., 2022). Our experiments were aimed to link phenotypic variation observed after storage under dry EPPO conditions with genetic markers to gain insights into potential candidate genes influencing seed longevity in rice.

### A pre-test showed the optimum aging duration for the GWAS panel

Individual seeds within a seed lot lose viability at different times in the form of a cumulative normal distribution of negative slope, i.e., the survival curve (Ellis and Roberts, 1980). The rate of viability loss over time, when transformed on a probit scale, exhibits a linear relationship, and forms the basis of the viability equations (Ellis and Roberts, 1980). Since deriving longevity parameters based on survival curves for all 300 accessions is laborious and would have required too many seeds from the genebank, we firstly chose to age seed samples of 20 rice accessions selected from different GWAS population groups (pre-test) under dry-EPPO conditions. The pre-test confirmed previous results (Prasad C. T. et al., 2022) that loss in viability under EPPO conditions was relatively fast compared with the ambient control and pressure control (EPPN) treatments at equivalent RH and temperature (Figure 1). Samples from all 20 accessions completely lost germination after 56 days EPPO aging, while hardly any significant loss in germination was noticed in that period under ambient storage. This confirmed the detrimental effect of oxygen on seeds during dry storage, which has been described for several crops (Roberts, 1961; Abdalla and Roberts, 1968; Justice and Bass, 1978; Ellis and Hong, 2007; Schwember and Bradford, 2011; Gerna et al., 2022). Based on the variation for survival in this pre-test (Figure 1), a 21-day EPPO aging period was selected as appropriate for phenotyping the larger GWAS population for germination parameters after aging.

We recorded a small negative response by certain accessions for the percentage of total seed germination in the pressure control treatment (EPPN), which increased with the duration of the treatment (Figure 1). Although the sensitivity is far less compared with the EPPO treatment, it is significant, and some genomic regions influence the sensitivity to EPPN 21 DOS (Supplemental Table 7). Further, the two isogenic lines used in the study of the *Rc* gene differed in their response to EPPN (Supplemental Figure 15B). Only the seeds from the line with a light-colored pericarp showed a delay in the rate of germination (increased t50), although this effect did not increase upon prolonged EPPN treatment. Yet, we do not know the mechanism of EPPN effects, to which the functional *Rc* gene seems to provide some tolerance. The duration-independent effect can be physical deterioration during pressure build-up or pressure decline. For accessions where the sensitivity also has a duration component, it seems more of a direct or indirect physiological or biochemical effect. A small deteriorating effect with EPPN aging was previously reported for seeds from some accessions in a diverse barley panel (Nagel et al., 2016).

### Diverse rice accessions show large variation for seed germination parameters under dry EPPO aging conditions

Although the 300 rice accessions used in the extended storage experiment were not selected based on prior knowledge of potential differences for seed longevity, they showed large variations for all the measured germination parameters after EPPO aging when compared with the control aging treatments (Figure 2). As expected, we also recorded a small negative effect in the main test with the pressure control aging treatment (EPPN 21 DOS). The effect of the EPPO aging treatment (EPPO 21 DOS) was corrected for trait values obtained by the pressure control (EPPN) and was depicted as ΔEPPO 21 DOS. We observed a good correlation between the observed trait values and normal expected values under different aging treatments, indicated by the high “W” values (Supplemental Figure 6), with a continuous variation indicating control of the traits by multiple genes. Although there was some variation in initial viability between seeds of different production years in control treatments (Control 0 DOS and Control 21 DOS), in general, the frequencies of germinating seeds were high (Supplemental Figure 7). This indicates that the germination potential of most accessions was well maintained during genebank storage and transport, followed by the 18 days equilibration. Moreover, the percentage of total seed germination under EPPO conditions of 20 rice accessions from the pre-test and the main test was highly correlated (*r*=0.94; Supplemental Figure 8), indicating similar performance under two independent aging tests. Since the effect of production year on the germination performance was small, we assumed that most variation is caused by genetic polymorphism, which allows for a GWAS approach.

### GWAS identifies genetic regions for total seed germination across different aging treatments

Previous genetic analyses for seed longevity in rice, using bi-parental mapping populations, have identified genomic regions on different linkage groups, namely *qRGR-1, qRGR-3, qLG-7, qLG-9*, and *qSS11* (Miura et al., 2002; Sasaki et al., 2005; Zeng et al., 2006; Xue et al., 2008; Jiang et al., 2011; Hang et al., 2015; Lin et al., 2015; Dong et al., 2017; Yuan et al., 2019). Aging conditions in these studies included either moist Accelerated Aging (AA) or CD experimental aging or ‘natural’ laboratory bench conditions with variable levels of RH and temperature during storage (reviewed in Hay et al. (2019)). Some studies with other crops have identified differences in QTLs when seeds of the same mapping populations were aged either under dry or moist conditions (Schwember and Bradford, 2010; Nagel et al., 2011; Nagel et al., 2016). Moreover, a weakness of identifying QTLs using a bi-parental mapping population, such as limited allelic diversity and lower mapping resolution as a result of the limited number of recombination events during the construction of a mapping population, can be overcome through GWAS (Korte and Farlow, 2013). Our GWAS identified a total of 12 unique regions (R1 to R12) on chromosomes 2, 4, 6, 7, and 8, and across different aging treatments (Figure 3 & Supplemental Table 7). Genomic regions ‘R7’ and ‘R9’ on chromosome 7 were common for non-corrected (EPPO 21DAS) and corrected (ΔEPPO 21DAS) aging treatments, which confirms the strong correlation between original and corrected trait values. Genetic analysis using the barley population found few overlaps between QTLs for tolerance to EPPN or EPPO aging treatments (Nagel et al. 2016), but common regions between the EPPN and EPPO treatments were not observed in our study woith rice seeds. GWAS for seed longevity parameters (*Ki*, -σ^-1^ and *p*50) in a large Indica rice panel identified major genomic regions on chromosomes 3, 4, 9, and 11 by storing seeds relatively dry (60% equilibrium RH and 45°C) (Lee et al., 2019). Our study did not return genomic regions previously identified in rice, possibly due to differences in the aging conditions, which can have a profound influence on the deterioration rate. Putative candidate genes identified in the genomic regions for control treatments relate more to having a role in the maintenance of seed quality during ambient dry storage, while those identified in EPPO treatment relate to tolerance against oxidation stimulated by elevated oxygen levels (Supplemental Table 8).

### The Rc allele on chromosome 7 explains tolerance by rice seeds to dry EPPO aging conditions

Our GWAS identified one QTL on chromosome 7 that was responsible for around 34 % of phenotypic variation in the germination capacity of EPPO-aged seeds (ΔEPPO 21 DOS). The most significant SNP (Chr7: 6067855) is located within the *Rc* gene (Figure 4A), encoding a bHLH transcription factor regulating proanthocyanidin production in the seed pericarp (Sweeney et al., 2006). The functional role of the *Rc* gene was confirmed in a separate storage experiment using seeds produced under the same environmental conditions from a pair of isogenic lines. Isogenic line *SD7-1D* with a functional *Rc* gene produces red pericarp seeds with a higher accumulation of proanthocyanidins (Figure 5A) and shows an enhanced longevity phenotype when compared to seeds from the *SD7-1d* line with light-colored pericarp due to the lack of a functional *Rc* gene (Figure 5C, Supplemental Figures 15 & 16). Proanthocyanidins are major flavonoids and are known to be involved in a multitude of functions because of their strong antioxidant capacity (Dixon et al., 2005; Lepiniec et al., 2006; Rauf et al., 2019; Wu et al., 2021).

Previous studies have reported a strong association between the accumulation of antioxidants in seeds and longevity (Nagel et al., 2015; Lee et al., 2017; Lee et al., 2019; Yuan et al., 2019). Therefore, it would be worth investigating whether in rice, proanthocyanidins in the pericarp act as a physical barrier for oxygen diffusion and thereby aid in the protection of the embryo against oxidative damage.

Our headspace ethanol analyses (Supplemental Figure 22) showed that the higher sensitivity of rice seeds with a light-colored pericarp (from line *SD7-1d*) is accompanied by relatively more anaerobic respiration during imbibition after seven days of EPPO storage than the seeds with a red pericarp (from line *SD7-1D*). An increase in ethanol production during storage has also been observed during prolonged storage of cabbage seeds on the laboratory bench (Kodde et al., 2012; Buckley et al., 2013). The ethanol production is likely related to accumulating oxidative damage to the mitochondria, pushing at least part of the seed tissues towards anaerobic respiration.

Fresh harvested seeds from the isogenic line *SD7-1D* with the functional *Rc* gene also exhibited more dormancy (Figure 5B & Supplemental Figure 14), confirming the results of a previous study (Gu et al., 2011). The reported difference in the rate of dormancy release during laboratory bench storage was also observed during EPPO aging, albeit at a faster rate. The latter agrees with genetic studies on *Arabidopsis* seeds, where the EPPO method has been used to accelerate seed dormancy release and mimic dormancy release by dry after-ripening (Buijs et al., 2018), indicating a role of oxygen in the release of this type of dormancy. Also, in *Arabidopsis*, mutants with reduced levels of pigments in the seed coat show reduced dormancy and storability under natural aging at room temperature (Debeaujon et al., 2000). Pipatpongpinyo et al. (2020) have demonstrated that genes controlling seed dormancy in rice are also involved in the regulation of survival under (moist) soil seed bank conditions. It would be interesting to study the storage behavior of seeds from the two isogenic lines under moist aging assays.

In addition to the link between the *Rc* gene and dormancy (Gu et al., 2011), our study provides evidence for a novel role of *Rc* in seed survival after dry storage and hence can provide an advantage if included in breeding programs to improve seed vigor and longevity. Incorporation of the functional *Rc* gene in popular rice varieties will not affect consumers’ preference for white rice since the pericarp is removed during polishing. However, it would be an added value for consumers preferring unpolished rice because proanthocyanidin antioxidants in the pericarp offer additional health benefits (Ravichanthiran et al., 2018; Lee et al., 2019; Saleh et al., 2019). A functional *Rc* gene had no effect on the other agronomic and yield traits (Supplemental Figure 13).

### Other candidate genes that have a potential role in controlling seed longevity in rice

Several candidate genes related to enzymatic antioxidant activity have been reported to influence seed longevity in rice (Long et al., 2013; Gayen et al., 2014; Huang et al., 2014; Sasaki et al., 2015; Petla et al., 2016; Galland et al., 2017; Nisarga et al., 2017; Yuan et al., 2019). Such genes did not appear in our study, but this was not unexpected since antioxidant enzymes cannot function under the dry experimental storage conditions used in our seed aging study (Gerna et al., 2022). Next to the *OsRc* gene, our study identified other candidate genes with a potential role in seed longevity (Supplemental Table 7 & 8). Candidate genes identified here can have a role in promoting seed germination, like *Cytochrome P450* (LOC_OS06g30500.1) with oxidoreductase activity (Kim and Tsukaya, 2002) and *Expansins* (Chen and Bradford, 2000; Yan et al., 2014). Other candidate genes revealed under dry EPPO storage suggest that heat response proteins, protease inhibitor proteins, and proteins with DNA/RNA binding properties may play an important role in seed longevity. A previous GWAS of an Indica rice panel identified candidate genes related to seed longevity that are involved in DNA dependent transcription and repair of damaged DNA (Lee et al., 2019). *HIT4* (LOC_Os02g31960) a novel regulator involved in the heat-triggered reorganization of chromatin, exhibits properties like heat shock proteins (HSPs) (Wang et al., 2015). Small HSPs are known to offer a protective role in desiccation tolerance, membrane stabilization, and oxidative stress tolerance (Wehmeyer and Vierling, 2000). Chromatin reorganization during seed maturation is an important developmental aspect of seeds preparing for ex-planta survival (van Zanten et al., 2011). The protease inhibitor domain identified in our study (*LTPL-163/4*) occurs in proteins like lipid transfer proteins, alpha-amylase inhibitors, and seed storage proteins (SSPs). SSPs buffer the seed from oxidative stress and thus protect the other proteins required for seed germination and seedling growth (Nguyen et al., 2015). Our study suggests that proteins involved in DNA/RNA binding activity (zinc finger proteins (*OsCT2FP8*); *Rc* (bHLH transcription factor) targeting gene transcription and Pentatricopeptide Repeat proteins involved in multiple aspects of RNA metabolism) may also be related to seed longevity. The *Arabidopsis thaliana Homeobox25 (ATHB25)* gene encoding a homeobox transcription factor targeting GA during seed development suggests a potential novel mechanism of seed longevity (Bueso et al., 2014). Pentatricopeptide Repeat proteins play an important role in RNA metabolism in organelles, particularly in mitochondria helping to overcome biotic and abiotic stresses (Xing et al., 2018). Further research is needed to confirm if these genes indeed influence rice seed longevity under dry storage conditions.

### Concluding remarks

In this study, we explored the natural variation for seed survival by storing dry seeds of diverse rice germplasm under a relative fast experimental aging method involving elevated oxygen conditions (EPPO). Our study demonstrates the power of GWAS, as it enabled the identification of the *Rc* gene as having a major role in conferring tolerance of rice seeds against elevated oxygen levels during dry storage. Our validation experiment clearly established that seeds with a functional *Rc* gene, responsible for PA synthesis and accumulation in the pericarp, offered greater resistance to oxidative damage. Favorable haplotypes of the *Rc* gene could be exploited in marker-assisted breeding, genomic selection, and genetic engineering to improve seed vigor and longevity in rice and other crop species. Future work will be required to validate the potential role of other candidate genes identified in this study and their targets in rice seed longevity. Considering the confirmed negative effect of oxygen on seed survival during storage, we strongly advocate storing dry seeds under anoxic conditions to extend shelf-life and maintain seed quality, particularly in hot humid climates. This approach will have far-reaching positive implications on seed conservation and food and nutritional security.

## MATERIALS AND METHODS

### Seed materials

A total of 300 rice accessions selected from the 3K panel (Rellosa et al., 2014) were used. The accessions represent diversity within the Indica variety group consisting of improved and traditional varieties from 34 different countries representing 12 rice growing areas of three continents (Supplemental Figure 1). Seeds were received from the International Rice Genebank Collection (IRGC) at the International Rice Research Institute (IRRI), Philippines. Seed samples of these accessions were produced in different years, ranging from 2009 to 2016 (Supplemental Table 1). In contrast to normal practice for dispatching seed samples from the IRGC, the seed samples used in our study had not received a hot water treatment for the elimination of potential contamination with nematodes since the hot water treatment may interfere with the seed longevity assay.

Isogenic lines with a functional *Rc* gene (*SD7-1D*) or a mutated *rc* gene (*SD7-1d*), both in the EM93-1 background, were provided by Prof. Gu, South Dakota State University, Brookings, USA. The introgression rice line containing genomic segment *qSD7-1* with the *SD7-1D* allele exhibits an enhanced seed dormancy (Gu et al., 2011). Seeds from these isogenic lines were produced in the greenhouse facilities at Wageningen University and Research under short-day conditions from April to September 2018. Germinating seeds were transplanted to plug trays filled with hydrated coco-peat mixture (Jongkind-Grond BV, http://www.jongkind-substrates.com). The trays were placed in a climate chamber maintained at 28/20°C with 10 h light/14 h dark during a day/night cycle. The growing medium was kept moist by intermittently adding demineralized water or Hyponex nutrient solution (https://www.hyponex.co.jp/). Three healthy 25-day-old seedlings were transplanted into each of 15 plastic pots filled with coco-peat: soil mixture. Three pots (nine plants) for both isogenic lines formed one block, and five blocks were established. A spacing of 300 mm between the pots and 600 mm between blocks was maintained. Each plant was tagged on the day of first panicle emergence, and seeds were harvested 40 days later. Tagged panicles from each plant were harvested separately, air-dried in the greenhouse for three days, and cleaned to select fully mature seeds. Cleaned fresh seeds were dried at 30% RH, sealed in polyethylene-lined aluminum pouches, and stored at -28°C until they were used in experiments.

### Seed aging treatments

Experimental aging of rice seeds under elevated partial pressure of oxygen (EPPO) was carried out according to the protocol described by (Groot et al., 2012; Prasad C. T. et al., 2022) with slight modifications. The required number of seeds for each line was transferred to a paper bag (63 × 93 mm packet size, 60g/m2 bleached Kraft paper, Baumann Saatzuchtbedarf GmbH, Germany, https://www.baumann-saatzuchtbedarf.de). The paper bags were rolled and secured with adhesive tape.

The seed packets together with silica gel (120g per tank, held in a 15 Denier nylon stocking) were equilibrated to 40% RH at 20°C in a controlled humidity cabinet for 18 days. Equilibrated seeds and silica gel were placed inside steel tanks (12 L for GWAS and 1.5 L for other experiments), and the tanks were closed immediately. Care was taken to ensure that the seed material and silica gel were not left on the laboratory bench for more than two minutes. The purpose of including the RH-equilibrated silica gel was to buffer the RH inside the tank upon filling it with dry compressed air.

The steel tanks with seed samples were filled slowly (approximately 0.6 MPa per minute) with either compressed air or pure nitrogen gas until the pressure reached 20 MPa (200 bars). Filling the tank with compressed air to 20 MPa resulted in a 4.2 MPa partial pressure of oxygen (PO2), called EPPO treatment. Filling a tank already containing air at ambient pressure with pure nitrogen gas (Grade, Nitrogen 5.0; Purity, ≥99.999 %; Linde Gas, Schiedam, The Netherlands) to 20 MPa resulted in 0.021 MPa PO2 (as in ambient air) and 19.978 MPa PN2, this is called EPPN treatment. The EPPN tank was intended as a control treatment for potential high-pressure effects. For the ambient pressure control treatment, the seed packets and silica gel stockings were placed in 1.5 L airtight Kilner glass jars. The PO2 inside the glass jars will be 0.021 MPa since, at sea level, air with a total pressure of 0.1 MPa is a mixture of gases containing 78% nitrogen, 21% oxygen, and 1% other gases. EPPO and EPPN tanks were placed inside a tub filled with chilled water to reduce heat build-up during filling. After completing the filling process, the tanks were left on the laboratory bench (20°C) overnight. For the pre-test and GWAS, two replications of the treatments were executed; for other experiments, single treatments with four biological replicates were performed. Due to the relatively small genebank sample size, 40-45 seeds were used per replicate.

The filled EPPO, EPPN tanks, and ambient pressure glass jars were placed in an incubator set at 35°C. The temperature increase (20 to 35°C) results in a measured increase of the air RH from 40 to 43.5% (Prasad C. T. et al., 2022), while the 20 MPa pressure results in a further theoretical increase of equilibrium RH from 43.5 to 50% (Okos et al., 2019). At the end of each time point, the tank(s) and jars were taken from the incubator and the tank pressure was checked for any leakage during storage, which had not occurred. The tanks and jars were left on the laboratory bench to bring the temperature of the tank to room temperature (RT). The pressure in the tank(s) was slowly released with an average relative pressure decline at a maximum of 0.5% per minute by connecting them to a computerized flow control device. After the pressure release process and retrieval of the samples, the water activity of the silica gel was checked in the laboratory (i.e., at 20°C) to confirm that the RH inside the tank (at ambient temperature and pressure) was 40%. Water activity was measured using HC2-AW probe connected to a HygroLab 3 display unit (Rotronic Measurement Solutions, https://www.rotronic.com). The treated seed samples were stored in hermetically sealed aluminum pouches at -20°C until used for the germination test. Sealed aluminum packets from the freezer were warmed to room temperature before opening, and the seed packets were placed in the seed drying cabinet (at 30% RH and 20°C), for at least overnight, before the germination test was conducted.

Since, for a significant number of lines, we observed differences between 21 days ambient control and 21 days EPPN pressure control treatment, we corrected for all germination trait values in 21 days EPPO aging treatment by compensating for the difference observed between ambient control and pressure control, replication wise in each accession. The corrected EPPO values were depicted as ‘ΔEPPO’ (Prasad C. T. et al., 2022).

### Storage time points for aging treatments in different experiments

To determine the optimum duration for the EPPO aging treatment to screen the GWAS population, a pre-test was performed with a sub-set of 20 diverse rice accession randomly selected from among the 300 accessions. In the pre-test, the EPPO treatment had eight storage durations (7, 14, 21, 28, 35, 42, 49, and 56 days). EPPN pressure and ambient pressure control had four storage durations (3, 28, 42, and 56 days) apart from the initial control (0 days). The main experiment was established with seeds of all 300 rice accessions receiving EPPO treatment and control treatments for 21 days, including the initial control (0 days). To test the tolerance of seeds from the isogenic lines (*SD7-1D* and *SD7-1d*) to dry-EPPO aging, a separate aging experiment was performed. That EPPO treatment had thirteen storage durations (3, 7, 14, 21, 28, 35, 42, 49, 56, 63, 70, 77, and 84 days). Here the EPPN pressure and ambient pressure controls had four storage durations each (3, 28, 56, and 84 days) along with initial control (0 days).

### Phenotypic data

#### Seed germination parameters

Seed germination assays were essentially performed as described in the International Rules for Seed Testing (ISTA, 2018) with some modifications. Dry seeds were sown on two layers of dry filter paper (142 × 203 mm blue blotter paper, All Paper, Zevenaar, The Netherlands) placed in plastic trays (150 × 210 mm DBP Plastics, Antwerpen, Belgium). For the pre-test and GWAS experiments two replicates of 40-45 seeds each were used. In the germination assays for assessing dormancy, after-ripening, and longevity with seeds from the isogenic lines, four biological replicates with 45 seeds per replicate were used. Germination experiments were conceived as a randomized complete block design.

Germination was initiated by dispensing 50 ml of demineralized water to each tray. Watered trays were stacked along with a watered tray without seeds, one at the bottom and one at the top. The stack of watered germination trays (maximum of 18) was wrapped in transparent plastic bags to prevent moisture loss due to evaporation. The prepared stacks were incubated in a climate chamber maintained at 25°C and continuous dark conditions. Germination was followed daily for up to 14 days by making images at frequent intervals using a digital camera (Nikon D80, http://www.nikon.com). Seeds were considered germinated if the radicle protruded by at least 2mm. Germination parameters, namely, total seed germination on the 14^th^ day (GMAX, in percentage), time for 50% germination (t50, in h), uniformity of germination (U9010; time interval between 10 and 90% seed germination), area under the germination curve until 250 h (AUC(250)) and mean germination time (MGT, in h) were calculated by automatically scoring germination over time with GERMINATOR software (Joosen et al., 2010). Total normal seedlings (TNS, in percentage) was determined on the 7^th^ and 14^th^ day of the germination test. Normal seedlings were evaluated according to the ISTA Handbook on Seedling Evaluation (Don, 2006). Seedlings with an intact, healthy shoot, and root system were evaluated as normal. The data for total seed germination for GWAS were transformed to a probit scale, where 50% germination is equivalent to 0 probits and lower germination values are negative.

#### Seed after-ripening and dormancy-breaking treatment

To analyze the speed of after-ripening after harvesting, freshly harvested air-dried seeds from the two isogenic lines were stored in an incubator maintained at 30% RH and 20°C. Seed samples were taken at weekly intervals to assess the release of dormancy. For the control treatment, dormancy was broken using a dry heat treatment (7 days at 50°C) as described in International Rules for Seed Testing (ISTA, 2018).

#### Measuring agronomic traits

Plant architecture and yield components for the isogenic *SD7-1D* (*Rc*) and *SD7-1d* (*rc*) lines were measured from nine plants or hill in each block at harvest maturity. Plant height (in mm) was measured from the soil level to the tip of the tallest spikelet on the panicle. The number of tillers and reproductive tillers (tillers with panicles) per plant was counted manually. Panicle length was measured from the base of the spikelet to the tip of the latest spikelet on the panicle. Days to flower initiation were recorded when the first panicle on the plant became visible. Yield components like the number of spikelets and fertile spikelets per panicle from tagged panicles were counted using an automatic seed counter. Hundred seed weight (in g) was measured on pooled seeds as per the procedure described in the International Rules for Seed Testing (ISTA, 2018). For weight determination, eight replicates of 100 seeds equilibrated at 30% RH for 14 days were used. Other seed morphological traits like seed length and width (in mm) on four replicates of 96 seeds from pooled seed samples from each isogenic line were measured using VideometerLab Instrument (Videometer A/S, Denmark, https://videometer.com/).

#### Analysis of mitochondrial quality

An ethanol assay for mitochondrial integrity was performed following the procedure described in Kodde et al. (2012) with slight modifications. Dehulled seeds were equilibrated at 30% RH at 20°C for four days. Five hundred milligrams of seeds were placed in 20 ml clean glass vials, and a calculated amount of Milli-Q water was added to the vials to achieve 25% seed moisture content. The vials were immediately sealed with aluminum crimp caps lined with rubber to prevent leakage and central septa to draw headspace air sample. The vials containing moistened seeds were placed in a pre-heated incubator at 50°C. Ethanol concentration was measured 24 hours after incubation by sampling 0.3 ml from the headspace with a modified breath analyzer (Alcotest 6810, Drägerwerk AG&Co. KGaA, Germany, https://www.draeger.com). Sample measurement outside the incubator was performed within 5 seconds.

#### Population structure, genetic diversity, and linkage disequilibrium decay

The population structure of the 300 rice accessions used in the study was analyzed using ADMIXTURE software (Alexander et al., 2009) with 3K-RG 1 million (M) GWAS SNP Dataset (release 1.0, http://snp-seek.irri.org/_download.zul). ADMIXTURE was run on the filtered SNP dataset with *K* ranging from 2 to

10. The optimum *K* value was selected based on ADMIXTURE’s cross-validation (CV) procedure. The *Q*-matrices from the ADMIXTURE analysis were visualized in R package *‘pophelper’* (Francis, 2017). Principal component analysis (PCA) to summarize the major patterns of SNP variation in filtered SNP dataset was performed in ‘TASSEL v 5.2.51’ (Kroon et al., 2007). The linkage disequilibrium (LD) decay measured based on the allele frequency correlation coefficients (*r*^*2*^) for all pairs of SNPs within 1000 Kb distance was computed using ‘*PopLDdecay*’ v3.40 program (Xu et al., 2018) with the following parameters: –MaxDist 1000 –MAF 0.05 –Het 0.4 –Miss 0.99. The median value of *r*^*2*^ in each bin for all chromosomes was then averaged to produce a final *r*^*2*^ estimate for a bin. LD decay rate was measured as the chromosomal distance at which the average *r*^2^ dropped to half its maximum value. The results of PCA were visualized in R Package ‘ggpubr’ and LD statistics were visualized in R Package ‘ggplot2’.

#### Genome-wide association analysis

The genome-wide association study was run using 3K-RG 1M GWAS SNP Dataset available at the Rice SNP-seek database (Mansueto et al., 2017). The selection and sequencing of rice accessions in the 3K-Rice Genome Project have been described previously (Rellosa et al., 2014). First, the downloaded SNP data was handled in PLINK (Purcell et al., 2007; Chang et al., 2015) to prepare input files for subsequent analysis. TASSEL v5.2.51 (Kroon et al., 2007) was used to filter SNP data for the rice population used in this study. Finally, the 1M SNP dataset was filtered to retain SNP markers with MAF ≥ 5% resulting in 128,667 biallelic markers. The density of these filtered SNPs across the chromosomes is illustrated in Supplemental Figure 2. Association analysis was run using a compressed mixed-linear model (CMLM; Zhang et al. (2010)), which accounts for population structure and family kinship (relatedness) implemented in the R Package ‘GAPIT version 2’ (Tang et al., 2016). The underlying regression model for association mapping analysis is:

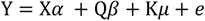

Where Y is the vector for phenotypic values, X represents the vector of genotypes at the candidate marker, *α* is the fixed effect for the candidate marker, Q is a matrix of principal components, *β* is the fixed effect for population structure, K is the relative kinship matrix, *μ* is a vector of random effects related to family identity, and *e* represents residual effects. Including Q and K matrices in the model helps to reduce the spurious false positives. The degree of correlation with population structure varies between traits. Therefore, GAPIT includes the optimal number of PCs to be included in the model based on the forward model selection method using the Bayesian Information Criterion (BIC). The genome-wide significant thresholds in this GWAS used a suggestive upper threshold determined using Bonferroni correction at *α* = 0.05 of -log_10_(0.05/128667) = 6.41 for extremely significant associations and lower suggestive threshold *p*-value of -log_10_(*p*) = suggestive upper threshold – 2 = 4.41 for significant associations.

#### Candidate gene selection

The upper limit of the LD decay rate is ∼500 Kb in rice (Mather et al., 2007). Since the average LD decay rate estimated in this population is ∼200 Kb, we applied a 200 Kb window on either side of the significant SNP (total ∼0.4 Mb) to investigate the local LD pattern and search for putative candidate genes (Huang et al., 2010). LD analysis was performed in Haploview 4.2 program (Barrett et al., 2005) to calculate LD structure and visualize the discrete haplotype block in the ∼0.4 Mb region harboring significant SNP. Gene models in the candidate regions and their known annotations were obtained from the MSU Rice Genome Database (Release 7; http://rice.plantbiology.msu.edu/cgi-bin/gbrowse/rice/). Promising candidate genes classified based on gene ontology and only those sufficiently described with relevant information from the closest *Arabidopsis* orthologs were selected. Genes described as transposons and retrotransposons were not considered.

#### Multiple sequence alignment

The full-length coding sequence for the *Rc* gene (LOC_Os07g11020) for *Oryza rufipogon* (IRGC 105491) was obtained from NCBI (Sweeney et al., 2006) and for *Oryza sativa* cv. MUTTU SAMBA::IRGC 36333-1 (IRGC 121441) & CT 9737-6-1-1-2-2P-M::IRGC 117330-1 (IRGC 120922) were downloaded from the Rice SNP-Seek Database (http://snp-seek.irri.org/_download.zul). Nucleotide sequences were translated into amino acids using the ‘transeq’ program implemented in EMBOSS package (Rice et al., 2000). The DNA and protein sequences were aligned using the ‘CLUSTALX 2.1’ program (Wilm et al., 2007).

#### Statistical analysis and data visualization

Germination parameters calculated as the mean of two separate germination tests were used for pre-test and association analysis. Germination parameters for other experiments are expressed as the mean value of four biological replicates. Tolerance for variation in germination percentages between two replicates was applied as per the tolerance tables given in the International Rules for Seed Testing (ISTA, 2018). Descriptive statistics, statistical analysis, and graphing were conducted in OriginPro 2022 (https://www.originlab.com/). Phenotypic data with percentage values were probit-transformed to improve normality. Two-way ANOVA was applied to observe genotype-treatment and its interaction effects on different traits. Paired t-test was used to compare the means of different traits for contrasting genotypes and treatments. Pairwise Pearson’s correlation coefficient was estimated to compare different traits. The effect of storage on dormancy breaking (after-ripening) followed by loss of viability of the two isogenic lines was modeled by probit analysis (Whitehouse et al., 2015). For all experiments, significance was determined by *p*<0.05 (\**p*<0.05, \*\**p*<0.01, \*\*\**p*<0.001). Box plots represents standard box setting with first and third quartile split by the median and the whiskers extend to a maximum of 1.5x interquartile range beyond the box. Principal component analysis (PCA) on phenotypic data, Genome-Wide Association Analysis, and data visualization were performed in R statistical program v3.5.0 (R Development Core Team, 2021). Manhattan plots and QQ plots to visualize GWAS results were executed using the R package ‘CMplot’ (Yin et al., 2021). PCA on germination parameters and visualization was performed using R packages ‘FactoMineR’ and ‘factoextra’ (Lê et al., 2008). VideometerLab software v1.8 (Videometer A/S, Denmark, https://videometer.com/) was used to extract seed morphological data of isogenic lines from the images.

## ABBREVIATIONS

AA: Accelerated aging test
AE: Allelic affect
ANOVA: Analysis of variance
AUC(250): Area under the germination curve until 250 h
aw: Water activity
bHLH: Basic helix-loop-helix transcription factors
BIC: Bayesian information criterion
bp: base pairs
CD: Controlled deterioration
CMLM: Compressed mixed linear model
DOS: Days of storage
DAI: Days after imbibition
EPPO: Elevated partial pressure of oxygen
EPPN: Elevated partial pressure of Nitrogen
ERH: equilibrium relative humidity
GAPIT: Genome Association and Prediction Integrated Tool
g: grams
GMAX: Maximum germination
GWAS: Genome-wide association study
h: hour
IRGC: International Rice Genebank Collection
IRIS: International Rice Information System
IRRI: International Rice Research Institute
*K*i: Initial viability in normal equivalent deviates
Kb: kilo bases
L: litre
LD: Linkage disequilibrium
LEA: Late embryogenesis abundant proteins
MAF: Minor allele frequency
M: million
Mb: Mega bases
ml: millilitre
MGT: Mean germination time
mm: millimetres
NED: Normal equivalent deviates
*P*50: Length of time for viability to fall to 50%
PA: Proanthocyanidin
PCA: Principal Component Analysis
PO2: Partial pressure of oxygen
QTL: Quantitative trait loci
RFO: Raffinose family oligosaccharides
RNS: Reactive nitrogen species
ROS: Reactive oxygen species
RT: Room temperature
Sigma (-σ^-1^): Length of time for viability to fall by 1 NED
SNP: Single nucleotide polymorphism
TF: Transcription factors
TNS: Total normal seedlings
t50: Time to reach 50% of the maximum germination
U9010: Uniformity of germination

## ACKNOWLEDGEMENTS

We would like to thank Prof. Xing-You Gu, Agronomy, Horticulture and Plant Science Department, South Dakota State University, Brookings, SD, USA for providing the isogenic lines *SD7-1D* and *SD7-1d*. Narendra Kumar Kata is acknowledged for contributions in conducting the germination tests. The authors thank Dr Wilma van Esse and Dr Niteen Kadam (Wageningen UR, The Netherlands) for constructive inputs during the research. We are grateful to Dr Vijay Dunna (ICAR-Indian Agricultural Research Institute, New Delhi) for critical reading of the manuscript.

## AUTHOR CONTRIBUTIONS

MP, GA, FH, KLM and SG designed the research; MP and JK performed the experiments and data analyses; MP, GA, FH and SG wrote the paper.

## Supporting Information

**Supplemental Table 1**. List of germplasm/accessions used in the study with details.

**Supplemental Table 2**. Descriptive statistics and ANOVA results for total seed germination (GMAX, %) of all the accessions used in pre-test to determine the optimum storage time for the treatments in the main test.

**Supplemental Table 3**. Descriptive statistics and ANOVA results for germination traits across aging treatments of 300 diverse rice accessions from main test.

**Supplemental Table 4**. Phenotypic data of all the accessions used in association mapping.

**Supplemental Table 5**. Principal components based on Principal Component Analysis (PCA) on different germination parameters.

**Supplemental Table 6**. Results of genome wide association study (-log_10_(*p*) value) for total seed germination across aging treatments.

**Supplemental Table 7**. List of significant genomic regions and their details identified by genome wide association analysis for total seed germination across aging treatments.

**Supplemental Table 8**. List of candidate genes located within the linkage disequilibrium block harboring the significant SNP identified in genome wide association study.

## Legends

**Supplemental Figure 1** | **Geographic origin and distribution of 300 rice accessions used in the genome-wide association study**. Geographic origin of rice accessions. Colors on the world map corresponds to the number of genotypes from the origin country.

**Supplemental Figure 2** | **Genome-wide SNP density**. The plot was generated using filtered SNPs (MAF ≥ 0.05) from 1M GWAS SNP dataset and indicate SNP density for a bin size of 1Mb on each chromosome. The color intensity corresponds to the number of polymorphic SNPs within 1Mb bin size according to the legend at right.

**Supplemental Figure 3** | **Variation for percentage total seed germination of 300 rice accessions under EPPO aging treatment. A**. Rice accessions are sorted on x-axis based on their total seed germination values, from high (left) to low (right). **B**. Representative germination images of rice accessions showing variable degree of tolerance to EPPO aging treatment compared with the controls. Scale bars in the bottom left corner of all the images indicates 50mm.

**Supplemental Figure 4** | **Phenotypic variation observed among 300 rice accessions for AUC(250) (A) and total normal seedlings (B)**. Aging treatments includes Control 0 days of storage (DOS) (■), Control 21 DOS (●), EPPN 21 DOS (▲), EPPO 21 DOS (♦) and ΔEPPO 21 DOS (+). Aging treatment was performed on seeds at 50% RH and 35°C for 21days (n= 2×40-45 seeds).

**Supplemental Figure 5** | **Seedling categories obtained from germination test of seeds stored under different aging treatments**. Aging treatments includes Control 0 days of storage (DOS); Control 21 DOS; EPPN 21 DOS, EPPO 21 DOS and ΔEPPO 21 DOS. The value under each seedling category in the bar chart indicate the average values of two germination replication with 40-45 seeds per replicate.

**Supplemental Figure 6** | **Phenotypic variation observed as frequency distribution among 300 rice accessions for total seed germination in the main-test**. Histogram with normal distribution curve overlay showing variation for percentage total seed germination under different aging treatments. Aging treatments includes Control 0 days of storage (DOS); Control 21 DOS; EPPN 21 DOS, EPPO 21 DOS and ΔEPPO 21 DOS. Under each germination parameter comparison is made between original values (row-1) and transformed values (row-2). Data was probit-transformed and normality statistics “W” indicates the correlation between observed values and normal expected values.

**Supplemental Figure 7** | **Variation for total seed germination under different aging treatments for rice accessions produced in different years. A**. Number of rice accessions representing different year of seed multiplication. **B**. Box plot showing distribution and comparison of means over different year of seed multiplication under different aging treatments for percentage total seed germination. Aging treatments includes Control 0 days of storage (DOS); Control 21 DOS; EPPN 21 DOS, EPPO 21 DOS and ΔEPPO 21 DOS. Each dot in the box plots represents average value from two germination test with 40-45 seeds per replicate.

**Supplemental Figure 8** | **Pearson’s pairwise correlation plot for percentage total seed germination**. Correlation plot between pre-test and main-test for total seed germination under 21 days EPPO aging treatment for 20 rice accessions. The germination values indicate average values of two germination replication with 40-45 seeds per replicate in both pre- and main-test.

**Supplemental Figure 9** | **Results of Admixture analysis. A**. Cross-validation plot for the 1M SNP dataset used in the admixture analysis. **B**. ADMIXTURE analysis for K = 2 & 5 indicating sub-populations.

**Supplemental Figure 10** | **Population structure and genetic diversity of the rice accessions used in the genome-wide association study. A**. Principal component analysis (PCA) indicating genetic variation based on filtered SNPs in the rice population used. Left PCA plot - first vs second PCs and right PCA plot - second vs third PCs. **B**. Scree plot showing the Eigen values and cumulative variability for each principal component on the filtered SNPs across 300 rice accessions. **C**. Number of lines representing different rice sub-populations. **D**. Genome-wide LD decay for average distance between SNP markers in the rice population.

**Supplemental Figure 11** | **Protein sequence of *Rc* protein generated with coding sequence in ClustalX 2.1**. Protein sequence comparison of *Rc* (*Oryza rufipogon* (IRGC 105491) and *O. sativa* cv. MUTTU SAMBA::IRGC 36333-1 (IRGC 121441)) and *rc* (*O. sativa* cv. CT 9737-6-1-1-2-2P-M::IRGC 117330-1 (IRGC 120922)). The 14 bp deletion in exon 7 in cv. CT 9737-6-1-1-2-2P-M::IRGC 117330-1 (IRGC 120922) results in premature stop codon at position 474 (indicated by * and highlighted in yellow color) resulting in light colored pericarp.

**Supplemental Figure 12** | **Allelic variation for *Rc* alleles among WT (SS18-2) and isogenic lines (*SD7-1D* and *SD7-1d*)**. Uppercase and lowercase letter indicate the mutation in exons and introns, respectively. Sequence in red color indicates sequence from weedy rice line SS18-2. Isogenic line with dormancy promoting allele *SD7-1D* have functional *Rc* gene responsible for seed red-color pericarp and dormancy decreasing allele *SD7-1d* have mutated *rc* gene responsible for seed with light-color pericarp.

**Supplemental Figure 13** | **Comparison of plant growth and seed traits in the isogenic lines *SD7-1D* and *SD7-1d*. A**. Whole plant, **B**. Plant height (mm), **C**. Number of productive tillers per plant, **D**. Number of seeds per panicle, **E**. 100 seed weight (g), **F**. Seed length (mm), and **E**. Seed width (mm). Isogenic line with dormancy promoting allele *SD7-1D* have functional *Rc* gene responsible for seed red-color pericarp and dormancy decreasing allele *SD7-1d* have mutated *rc* gene responsible for seed with light-color pericarp.

**Supplemental Figure 14** | **Total seed germination of seeds 3 days after harvest (DAH) and after dormancy breaking treatment (DT; heat treatment at 50°C for 7days) between isogenic lines *SD7-1D* and *SD7-1d***. Germination values indicate average values with four biological replicates of 40 seeds each at 14 days after imbibition. Isogenic line with dormancy promoting allele *SD7-1D* have functional *Rc* gene responsible for seed red-color pericarp and dormancy decreasing allele *SD7-1d* have mutated *rc* gene responsible for seed with light-color pericarp.

**Supplemental Figure 15** | **Comparison of seed germination parameters of isogenic lines *SD7-1D* and *SD7-1d* under ambient, EPPN and EPPO aging treatments. A**. Percentage total seed germination and **B**. time to reach 50% germination (t50, in h). Freshly harvested seeds equilibrated to 40% RH at 20°C for 22 days were used. Aging treatment was performed at 50% RH and 35°C for different storage periods. Each dot in the plot indicates average values of germination test evaluated on four biological replicates of 45 seeds each at 14 days after imbibition (DAI).

**Supplemental Figure 16** | **Comparison of seed germination of isogenic lines *SD7-1D* and *SD7-1d* under ambient, EPPN and EPPO aging treatments**. Representative germination images on the first, second, third and fourth row show germination from seeds subjected to different aging treatments for 3 days of storage (DOS), 28 DOS, 56 DOS and 84 DOS, respectively. Images tagged at the lower right bottom with green color represents germination at 5 days after imbibition (DAI), with red color represents germination at 7DAI and with orange color represents germination at 14DAI. Scale bar in all the images at the lower left bottom indicate 50mm.

**Supplemental Figure 17** | **Effect of different seed aging treatments on ethanol production**. Headspace ethanol levels in isogenic lines *SD7-1D* and *SD7-1d* after 7 days of storage (DOS) **(A)**, 14 DOS **(B)** and 21 DOS **(C)** under different aging treatments. Aging treatment was performed with seeds at 50% RH and 35°C, kept under ambient control, pressure control (EPPN) and high oxygen (EPPO) for different storage durations (7 DOS, 14 DOS and 21 DOS). Headspace ethanol was measured from 100mg seed samples at 25% moisture in 20ml glass vials incubated for 24 h at 50°C. Bar graphs shown are mean ± SE of mean of two biological replicates. Different letters over the bars indicate significant differences between the aging treatments (*p* < 0.05).

